# Human Argonaute2 and Argonaute3 are catalytically activated by different lengths of guide RNA

**DOI:** 10.1101/2020.07.16.207720

**Authors:** Mi Seul Park, GeunYoung Sim, Audrey C. Kehling, Kotaro Nakanishi

## Abstract

RNA interfering is a eukaryote-specific gene silencing by 20∼23 nucleotide (nt) microRNAs and small interfering RNAs that recruit Argonaute proteins to complementary RNAs for degradation. In humans, Argonaute2 (AGO2) has been known as the only slicer while Argonaute3 (AGO3) barely cleaves RNAs. Therefore, the intrinsic slicing activity of AGO3 remains controversial and a long-standing question. Here, we report 14-nt 3′ end-shortened variants of let-7a, miR-27a, and specific miR-17-92 families that make AGO3 an extremely competent slicer by an ∼ 82-fold increase in target cleavage. These RNAs, named cleavage-inducing tiny guide RNAs (cityRNAs), conversely lower the activity of AGO2, demonstrating that AGO2 and AGO3 have different optimum guide lengths for target cleavage. Our study sheds light on the role of tiny guide RNAs.

## Introduction

MicroRNAs (miRNAs) are small non-coding RNAs that control gene expression post-transcriptionally (1, 2). Their sequences differ, but their lengths fall within a range of 20∼23 nt because the precursor miRNAs are processed by Dicer, which is a molecular ruler that generates size-specific miRNA duplexes (3, 4). After those duplexes are loaded into AGOs, one of the two strands is ejected while the remaining strand (guide strand) and the AGO form the RNA-induced silencing complex (RISC) (5). Therefore, the 20∼23-nt length is the hallmark of intact miRNAs. This size definition has been exploited as the rationale for eliminating ∼18-nt RNAs when AGO-bound miRNAs are analyzed by next-generation RNA sequencing (RNAseq). However, RNAseq without a size exclusion reported a substantial number of ∼18-nt RNAs bound to AGOs (6-8). Such tiny guide RNAs (tyRNAs) are abundant in extracellular vesicles of plants (9), but little is known about their roles or biogenesis pathways. In mammals, the roles of tyRNAs are even more enigmatic.

The goal of this study was to understand target-RNA cleavage by human AGOs. In 2004, two groups reported that only AGO2 showed the guide-dependent target cleavage in vitro (10, 11). Since then, AGO1, AGO3, and AGO4 were thought to be deficient in RNA cleavage, even though AGO3 shares the same catalytic tetrad with AGO2. Recently, we revealed that specific miRNAs such as 23-nt miR-20a make AGO3 a slicer, but the activity was much lower than that of AGO2 (12). Therefore, it remained unclear whether AGO3 becomes a highly competent slicer as well. We revisited this long-standing question by investigating the effect of the guide length on target cleavage and discovered the unexpected role of tyRNAs in the catalytic activation of AGO3.

## Results

Purified recombinant proteins of AGO2 and AGO3 (12) were pre-incubated with either of 8, 10, 12, 13, 14, 15, 16, or 23-nt single-stranded synthetic miR-20a whose 3′ 7∼15 nt are deleted, followed by addition of a cap-labeled target RNA (Fig. 1A) as previously reported (13). While AGO2 reduced slicing activity with a shorter guide, AGO3 showed the highest cleavage activity with the 14-nt guide (Fig. 1B top and middle). Notably, the slicing activity of AGO3 with the 14-nt miR-20a was about 30-fold higher than its 23-nt intact form (Fig. 1B bottom), which resulted in AGO3 being a comparative slicer to AGO2. Supporting this, the kinetics of target cleavage with the 14- and 23-nt guide showed opposite trends between AGO2 and AGO3 (Fig. 1C). These results suggest that AGO2 and AGO3 have different optimum lengths of guide RNA for target cleavage.

**Fig. 1.**
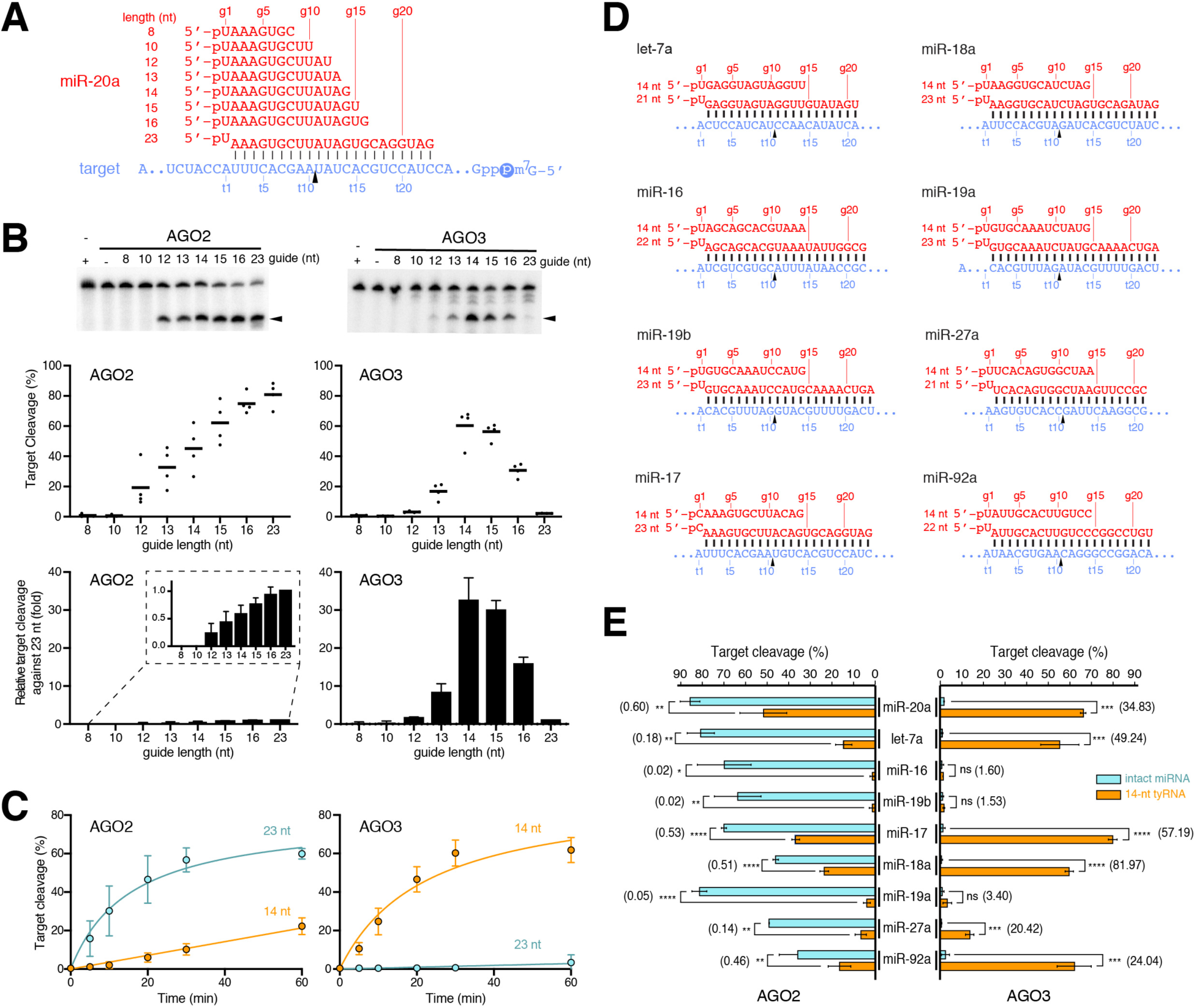
14-nt miR-20a brings out the slicing activity of AGO3. (A) Guide (red) and target (blue) RNAs used in B-C. The 60-nt target RNAs were cap-labeled. (B) In vitro cleavage assay by AGO2 and AGO3 with different lengths of miR-20a. Top: representative gel images. Middle: target cleavage percentages. Data are shown as mean (bar) and individual biological replicates (dots). Bottom: relative target cleavage (fold) with each guide length against 23 nt. Data are shown as Mean ± SD. (C) Time-course assay of AGO2 and AGO3 with the 14- or 23-nt miR-20a. Data are shown as Mean ± SD. (D) Guide and target RNAs used in E. (E) In vitro target cleavage using AGO2 and AGO3 with the intact miRNAs (cyan) or 14-nt (orange) tyRNAs. The number in parentheses indicates the relative target cleavage (fold increase) of each 14-nt tyRNA against the cognate intact miRNA. Data are shown as Mean ± SD. \**P* < 0.05; \*\**P* < 0.01; *** *P* < 0.001; **** *P* < 0.0001; ns, not significant (Student’s t test).

Intact miRNAs of let-7a, miR-16, and miR-19b are known to activate AGO2 but not AGO3 (12). To test whether their 14-nt tyRNAs serve as cityRNAs, recombinant AGO2 and AGO3 were programmed with either of their intact miRNA or tyRNA and subsequently incubated with the cap-labeled target RNA (Fig. 1D). Loading of the tyRNAs drastically decreased or ruined the slicing activity of AGO2, compared to that of their intact form (Fig. 1E). In contrast, not the 14-nt miR-16 or miR-19b, but the 14-nt let-7a conferred extremely competent slicing activity on AGO3. To find more cityRNAs, 14-nt tyRNAs of miR-17, miR-18a, miR-19a, miR-27a, and miR-92a (Fig. 1D) were tested for in vitro target cleavage. Again, AGO2 reduced slicing activity with their 14-nt tyRNAs whereas AGO3 became a remarkably competent slicer when loaded with all except for the 14-nt miR-19a (Fig. 1E). These results indicate that some cityRNAs make AGO3 a superior slicer to AGO2.

Unlike miRNA duplexes, 14-nt RNAs are too short to form stable double-stranded RNAs. Thus, we thought that such short RNAs could be loaded as a single-stranded RNA into AGOs. To test this idea, we performed a RISC maturation assay (14, 15). In the positive control experiment, the 23-nt siRNA-like duplex of miR-20a (p23ds) was used instead of p14ss (Fig. 2A). As expected, the provided 23-nt duplex was detected as its single intact strand in both AGO2 and AGO3 (Fig. 2B). Similarly, the intact 14-nt miR-20a was detected from both AGOs (Fig. 2B), demonstrating that AGO2 and AGO3 can incorporate the 14-nt ssRNAs in the cell lysate. Next, those assembled RISCs were immunopurified from the cell lysate and tested for slicing activity. FLAG-AGO2 cleaved RNAs very well when the lysate was incubated with the 23-nt siRNA-like duplex of miR-20a (23ds) (Figs. 2A and 2C). In contrast, FLAG-AGO3 became a very competent slicer when the 14-nt single-stranded miR-20a (14ss) was added to the lysate (Fig. 2C). To confirm that the observed RNA cleavage was due to the catalytic activity of AGO3, we repeated the experiment using a catalytically dead mutant, FLAG-AGO3 (E638A) (12). This mutant loaded both p14ss and p23ds to form the mature RISCs (Figs. 2A-B) but did not show slicing activity at all (Fig. 2C), proving that the catalytic center of AGO3 is essential for the cityRNA-dependent target cleavage. Lastly, we tested whether AGO3 incorporates 14-nt single-stranded RNAs within the cell to assemble a functional slicer. The 14-nt single-stranded miR-20a was modified, according to a previous report (16), to make it stable during and after transfection (14md in Fig. 2A). When programmed with the 14md, the recombinant AGO3 showed a slightly higher target cleavage than with the unmodified form (Fig. 2D), indicating that the modified guide retained the ability to catalytically activate AGO3. Then, HEK293T cells were co-transfected with a plasmid encoding FLAG-AGO2 or FLAG-AGO3 and either the 14ss, the 14md, or the 23ds. Immunopurified FLAG-AGO2 cleaved RNA very well with transfection of the 23ds (Fig. 2E). In contrast, FLAG-AGO3 cleaved the target RNA only when the 14md was co-transfected. These results demonstrate that AGO3 and 14-nt ssRNAs assemble into an active slicer in vivo.

**Fig. 2.**
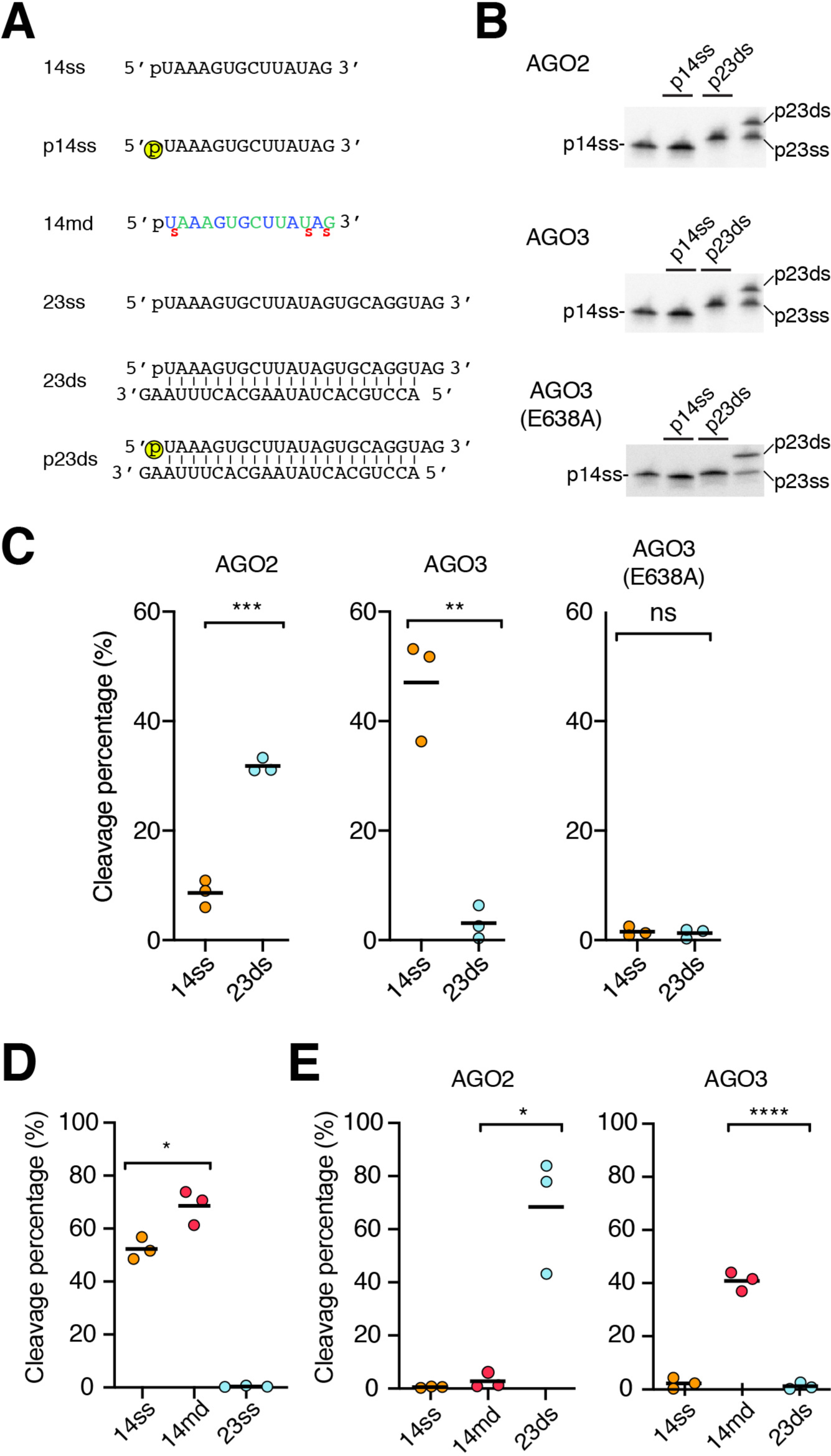
In vivo loading of 14-nt single-stranded guide RNA makes AGO3 a slicer. (A) 14- and 23-nt miR-20a variants. 14ss is identical to the 14-nt miR-20a in Fig. 1A. p14ss is identical to 14ss, except for the 5′-end radiolabeling (yellow circle). 14md is identical to 14ss, except for nucleotide modifications (blue: 2′-OMe, Green: 2′-F, red s: Phosphorothionate). 23ss is identical to the 23-nt miR-20a in Fig. 1A. 23ds is composed of a 23ss (top strand) and a passenger strand (bottom), the latter of which lacks a 5′ monophosphate group so AGOs load the 23ss. p23ds is identical to 23ds, except for the 5′-end radiolabeling. (B) RISC assembly assay. Either p14ss or p23ds was added to the lysate of HEK293T cells expressing FLAG-AGO2, -AGO3, or -AGO3 (E638A). (C) In vitro target cleavage using AGOs programmed in cell lysate. Either 14ss or 23ds was added to the lysate of HEK293T cells expressing FLAG-AGO2, -AGO3, or -AGO3 (E638A). (D) In vitro cleavage assay of recombinant AGO3 programmed with 14ss, 14md or 23ss. (E) In vitro cleavage assay using AGOs programmed within cell. Either 14ss, 14md or 23ds was transfected into HEK293T cells expressing FLAG-AGO2 or -AGO3.

## Discussion

AGO2 cleaves any RNAs including a sequence fully complementary to the guide RNA, which means that any guide RNAs can activate AGO2. This is not the case for the AGO3 activation. Only specific tyRNAs can serve as cityRNAs due to their unique sequences. These multiple requirements extremely limit the opportunities for catalytically activating AGO3. That is why the slicing activity has not been unveiled for a long time (10, 11). Since AGO3 has retained the catalytic center throughout its molecular evolution, the cityRNA-dependent slicing activity could have a conserved role in or beyond RNA interference when all the requirements are satisfied.

## Materials and Methods

### Cloning, expression, and purification of recombinant AGO proteins

Recombinant AGO2 and AGO3 were purified from the insect cells as previously reported (12, 14).

### In vitro cleavage assay

1 µM AGO proteins were incubated with 100 nM 5′ phosphorylated synthetic single-stranded guide RNAs for RISC assembly in 1× Reaction Buffer (25 mM HEPES-KOH pH 7.5, 5 mM MgCl_2_, 50 mM KCl, 5 mM DTT, 0.2 mM EDTA, 0.05 mg/mL BSA (Ambion), and 5 U/µL RiboLock RNase Inhibitor (Thermo Scientific)). 5′ cap-labeled target RNAs were added in the reaction for the target cleavage. The reaction was directly quenched with 2× urea quench dye (7 M urea, 1 mM EDTA, 0.05% (w/v) xylene cyanol, 0.05% (w/v) bromophenol blue, 10% (v/v) phenol). The cleavage products were resolved on a 7M urea 16% (w/v) polyacrylamide gel.

### Validation of modified 14-nt miR-20a by in vitro cleavage assay

1 µM recombinant AGO3 was incubated with the 14ss, the 14md, or the 23ss for 1 hour at 37 °C followed by target cleavage as described above.

### In vitro cleavage assay using FLAG-AGO programmed within the cell

10 µg of pCAGEN vector encoding FLAG-AGO2, FLAG-AGO3, or FLAG-AGO3 (E638A) was transfected into HEK293T cells. After 24 hours, the 14ss, the 14md, or the 23ds was transfected. 24 hours later, the cells were harvested and lysed by sonication. The amount of FLAG-AGO proteins in the cell lysate was normalized based on the western blot result. AGO was quantified by using a standard curve generated with known amounts of recombinant FLAG-AGO3 (14). The overexpressed FLAG-AGOs were immunoprecipitated with 50 μL anti-FLAG M2 beads. After the beads were washed with IP Wash Buffer, the cap-labeled 60-nt target RNAs were added for cleavage reaction.

## Acknowledgements

This work was supported by a Pelotonia Fellowship (to M.S.P.), a Center for RNA Biology Fellowship (to G.Y.S), the NIH (R01GM124320 and R01GM138997 to K.N.), and the Office of the Director, NIH (S10OD023582).

## Author Contributions

M.S.P., G.Y.S. and A.K. expressed and purified recombinant proteins. M.S.P and G.Y.S. performed biochemical assays. M.S.P., G.Y.S., and K.N. analyzed the data. K.N. designed the research and wrote the manuscript with input from the other authors.

